# Predictors of emergency department attendance following NHS 111 calls for children and young people: analysis of linked data

**DOI:** 10.1101/237750

**Authors:** Arne Wolters, Cicely Robinson, Dougal Hargreaves, Rebecca Pope, Ian Maconochie, Sarah Deeny, Adam Steventon

## Abstract

**Objectives:** To assess whether clinical input during calls to the NHS 111 telephone-based advice service is associated with lower rates of subsequent emergency department attendance and hospital admission.

**Design:** Although NHS 111 largely employs non-clinical call handling staff to triage calls using computerised clinical decision support software, some support is available from clinical supervisors, and additionally some calls are referred to out-of-hours General Practitioners (GP). We used linked data sets to examine GP and secondary care activity following calls to NHS 111, adjusting for the patient characteristics, signs and symptoms recorded during the NHS 111 call.

**Setting:** Out-of-hours care in three areas of North West London that have an integrated approach to delivering NHS 111 and out-of-hours GP care.

**Participants:** NHS 111 calls for children and young people aged 15 years or under. We excluded calls that were diverted to the emergency (‘999’) service or where patients were advised to go to an emergency department. This left callers who were either referred to a GP or advised to manage their health needs at home.

**Primary and secondary outcome measures:** The percentage of callers attending any emergency departments, major emergency department, or minor injury unit within ten hours of the NHS 111 call, and the percentage admitted to hospital following visits to emergency departments.

**Results:** Of the 10,356 callers, 2,898 (28.0%) were advised by NHS 111 to manage their health needs at home, with an appointment with an out-of-hours GP made for the remaining 7,458 (72.0%). 14.9% (432/2,898) of the callers who were advised by NHS 111 to manage their health needs at home attended an emergency department with ten hours, compared with 16% (1,207/7,458) of callers who had an out-of-hours appointment with an out-of-hours GP. After adjusting for patient characteristics, GP out-of-hours appointment was associated with lower rates of emergency department attendance (adjusted odds ratio, 0.86, 95% CI, 0.75-0.99),). When we subset emergency department types, a GP out-of-hours appointment was associated with lower rates of minor injury unit attendance (adjusted odds ratio, 0.32, 95% CI, 0.23 - 0.44) but not major emergency department attendance (adjusted odds ratio 1.06, 95% CI 0.90-1.24). There was no association with hospital admission. Review by an NHS 111 clinical supervisor was associated with fewer emergency department attendances (adjusted OR 0.77, 95% CI, 0.62-0.97).

**Conclusions:** Clinical input during or following out-of-hours calls to NHS 111 was associated with lower rates of emergency department utilisation for children and young people, though the reduction may be concentrated in lower intensity care settings. Thus, there may be potential to reduce the use of emergency care by providing access to clinical advice or out-of-hour services in other settings through the NHS 111 telephone service.

## INTRODUCTION

In light of increasing pressures on costs, health care systems are exploring approaches to labour substitution and workforce configuration.[1] For example telephone advice lines, are a common international approach [2] and used in the Netherlands[3], United States[4], Scotland[5], Australia,[6] and Norway [7]. One example from England is the main telephone-based service, NHS 111, which employs non-clinically trained staff to triage patients with urgent health care needs, usually outside of normal working hours. With the help of computerised clinical decision support software (‘NHS Pathways’), health advisors gather information on signs and symptoms to arrive at a decision, which might include dispatching an ambulance, advising the caller to attend an emergency department, advising the caller to visit an out-of-hours GP centre, or giving advice about how to manage the complaint at home without a face-to-face contact. Expanding the use of these telephone-based advice services is common strategy internationally to signpost patients to the most appropriate urgent and emergency care service and reduce the pressure on emergency rooms.[3–5,7]

NHS 111 is becoming the most common route for people to access out-of-hours primary care,[8] and it receives almost 15 million calls annually.[9] NHS commissioning guidance for England has recommended that NHS 111 form a “single entry point” for integrated care hubs that provide clinical assessment, advice and treatment, supported by referrals to existing primary, community, secondary and social care services.[10,11] However, there have been concerns that, by relying on non-clinically trained staff, NHS 111 might have an overly cautious approach to handling risk, leading to a greater use of emergency care compared with other approaches to triage out-of-hours care.[12,13] This concern was not abated by a controlled before-and-after evaluation in four pilot sites, which reported that the introduction of NHS 111 was associated with a 2.9% increase in emergency ambulance call-outs and no change in emergency department attendances in the pilot sites.[14] Furthermore, although 92% of callers reported being satisfied or quite satisfied with the service,[15] a quarter of parents calling the number did not fully have confidence and trust in the first call handler. [15] To address this updated commissioning guidance now includes recommendations for increased clinical input, [16] and NHS 111 calls now receive clinical input by a nurse or GP (clinical supervisor) in over 30% of cases,[17] and this policy is to be sustained.[18]

While a number of studies have assessed the impacts of telephone-based health care on the utilisation of emergency departments,[19–21] few studies have addressed the impact of clinical versus non-clinical workforce to triage requests for out-of-hours care on the ultimate destination of callers.[1,22] One study in Cambridgeshire placed GPs within a NHS 111 call centre, and arranged for the GPs to review the cases who would have otherwise been advised by health advisors to attend emergency departments.[23] Of the 1,474 cases reviewed, the GPs recommended emergency department attendance in only 27% of cases. However, this study did not have access to linked data sets, meaning that the implications for secondary care utilisation could not be assessed. For the current study, we linked primary and secondary care data sets for three areas in North West London, which have an integrated approach to delivering NHS 111 and out-of-hours general practice care, with a single provider responsible for both services. We examined how calls to NHS 111 for children and young people were routed, and to what extent subsequent emergency department attendances depend on what clinical input the call received from a NHS 111 clinical supervisor or GP out-of-hours care.

## METHODS

We examined calls to NHS 111 for children and young people, and compared emergency department utilisation between the group of patients who were referred to an out-of-hours GP and those who were advised by NHS 111 to manage symptoms at home. Thus, we excluded NHS 111 calls that were diverted to the emergency (‘999’) service or where patients were advised to go to an emergency department without first being reviewed by a GP. We expected to find higher levels of health need amongst the patients who were reviewed by a GP than amongst those who were advised by NHS 111 to manage their health needs at home. We then compared how often these groups attended emergency departments, both minor injury units and emergency departments, following the NHS 111 calls, adjusting for the patient characteristics, signs and symptoms that were recorded on the NHS 111 database.

### Data sets

We examined three areas of North West London (Hammersmith and Fulham, Kensington and Chelsea and Westminster), in which NHS 111 is provided by London Central & West Unscheduled Care Collaborative (LCW), a GP-led not-for-profit organisation. As well as operating the NHS 111 line, LCW provides out-of-hours GP services and other GP-led services including urgent care services in a number of hospitals. In those three areas, NHS 111 health advisors do not have access to the general practice electronic medical record, but they can make an appointment for the patient to see an out-of-hours GP. The health advisors are also supported by clinical supervisors.

North East London Clinical Support Unit provided episode-level data from the operational databases used to manage NHS 111 calls and out-of-hours general practice contacts by LCW. These were linked to data on emergency department from the Secondary Uses Service, which is an administrative database that is closely related to the Hospital Episode Statistics, and contains all records of secondary care activity for patients who were residents of Hammersmith and Fulham, Kensington and Chelsea and Westminster, or were registered with a general practice in that area. Data were pseudonymised, with direct identifiers such as name and address removed and the NHS number replaced by patient reference number, which was then used to link the three datasets together.

We structured the data into ‘NHS 111 call episodes’, which include contacts with the health advisors and ‘warm transfers’ to the clinical supervisors who were based within the NHS 111 call centre. In some cases, NHS 111 called the patient back, for example if the original call was disconnected; those call-backs were included within the call episode. Then, we restricted our attention to call episodes made outside of office hours (defined as 9am to 5pm, Monday to Friday), and by or on behalf of people aged 15 or younger between 1 July 2013 and 28 February 2015. The age threshold was chosen for consistency with previous studies.[24] We excluded a small number of calls that were missing the information required to link those records with GP and emergency department data, and those that were diverted to 999 services or directed to an emergency department.

### Study endpoints and exposure variables

We examined visits to any emergency department, then minor injury units and specialist emergency departments considered separately, that occurred within ten hours of the start of the NHS 111 call. Although NHS 111 typically advised attendance within four hours, we examined visits over ten hours because in some instances patients might have first sought additional advice for the same complaint. In secondary analysis, we examined hospital admissions that occurred following an emergency department visit, which we identified based on the attendance disposal code in the emergency department data. Thus, we excluded direct, urgent admissions that did not come via the emergency department.[25]

For each of the NHS 111 call episodes, we determined whether the patient had an appointment recorded with an out-of-office GP within 90 minutes of the start of the NHS 111 call.

### Statistical analysis

We used logistic regression modelling to test the association between the likelihood of attending either type of emergency department following the NHS 111 call and whether the patient had an appointment recorded with an out-of-office GP. This model controlled for a range of patient characteristics: the age of the patient in complete years; the ethnicity of the caller; socioeconomic deprivation score (based on quintiles of the Index of Multiple Deprivation for the small area of residence); the nature of the health problem presented; the length of the call episode (measured in minutes from the beginning of the initial call to NHS 111 to the end of the last call-back); whether the caller was transferred to a clinical supervisor within NHS 111 call centre (a ‘warm transfer’); and whether NHS 111 called the patient back as part of the call episode.

To simplify the analyses, the 157 diagnostic codes used by NHS 111 were aggregated by two authors (DH, IM); this resulted in the following 13 categories: breathing difficulty, cough, cold or influenza; febrile illness; diarrhoea or vomiting; abdominal pain, constipation or rectal pain; rash or skin problem; injury, limb problem or burn; allergy; ear nose or throat problem; unwell infant; blisters; chest pain; eye problem; or other (one of 60 infrequently-occurring diagnostic codes). Table A1 in the supplementary material lists the diagnostic codes included in each of these groups. In addition, some calls were coded as having a pre-determined plan or specific pathway, while some others were requests for health and social care information.

In addition to the study covariates listed above, our logistic regression models included dummy variables for the patient’s registered general practice (‘fixed effects’). These were intended to control for a range of unobserved variables relating to the approach to managing patients within general practices. We did this because out-of-hours health care seeking behaviours might depend on the nature of care given within normal office hours. Also, some general practices might more actively direct patients to NHS 111 for out-of-hours care than other practices, and callers from those practices might present with NHS 111 with differing health needs than patients at other practices.

Statistical analyses were performed using Statistical Analysis Software (SAS Enterprise Guide, 7.1)[26].

#### Ethics

We did not require research ethics approval for this study since it involved retrospective analysis of pseudonymised data. The local Caldicott Guardian approved the study.

#### Data Sharing

This study used pseudonymised data collected and linked for this study. Under this agreement the authors do not have permission to share the dataset. Though data could be made available to researchers who applied to, and gained approval from appropriate local data holders and guardians.

#### Patient involvement

No patients were involved in setting the research question, in the outcome measures, in the design, in the implementation of the study, or in the interpretation and writing up of results. As study participants are anonymous the results cannot be disseminated to them, however they will be disseminated to relevant patient groups through the Health Foundation’s communication channels and relevant news media. As patients were not directly involved in the research, they have not been thanked in the contributorship statement/acknowledgements.

## RESULTS

NHS 111 received 18,112 calls from the three areas of North West London between July 2013 and February 2015. Of these, 3,633 calls were missing fields (NHS number) required for data linkage (20.1%), 2,083 calls were made during working hours (11.5%), 1,117 calls were routed to 999 (6.2%), and 923 calls were immediately advised to attend emergency departments (5.1%). This left 10,356 calls for our analysis, for 7,651 distinct patients registered at 118 general practices (a flow diagram of cohort selection is shown in Figure A1 in supplementary material).

Figure 1 shows the flow of patients through the NHS 111, GP and emergency department services. Of the 10,356 callers, 2,898 (28.0%) were advised care at home, with an appointment with an out-of-hours GP made for the remaining 7,458 (72.0%). Compared with the patients who were advised care at home, those who were referred to a GP were younger (mean age 3.1 *vs*. 3.4 years, see Table 1). They were more likely to have each of the 12 conditions, with the exception of eye problems and the ‘other’ category (see Table 1). Patients who were referred to a GP were also more likely to have a pre-determined management plan (15.1% vs. 4.8%). They were more likely to present at the weekend (55.9% *vs*. 39.5% of calls) and typically followed NHS 111 calls that were shorter (mean 10.1 *vs*. 16.3 minutes) and a lower proportion were passed to a clinical supervisor (6.0% *vs*. 16.4%).

**Figure 1.**
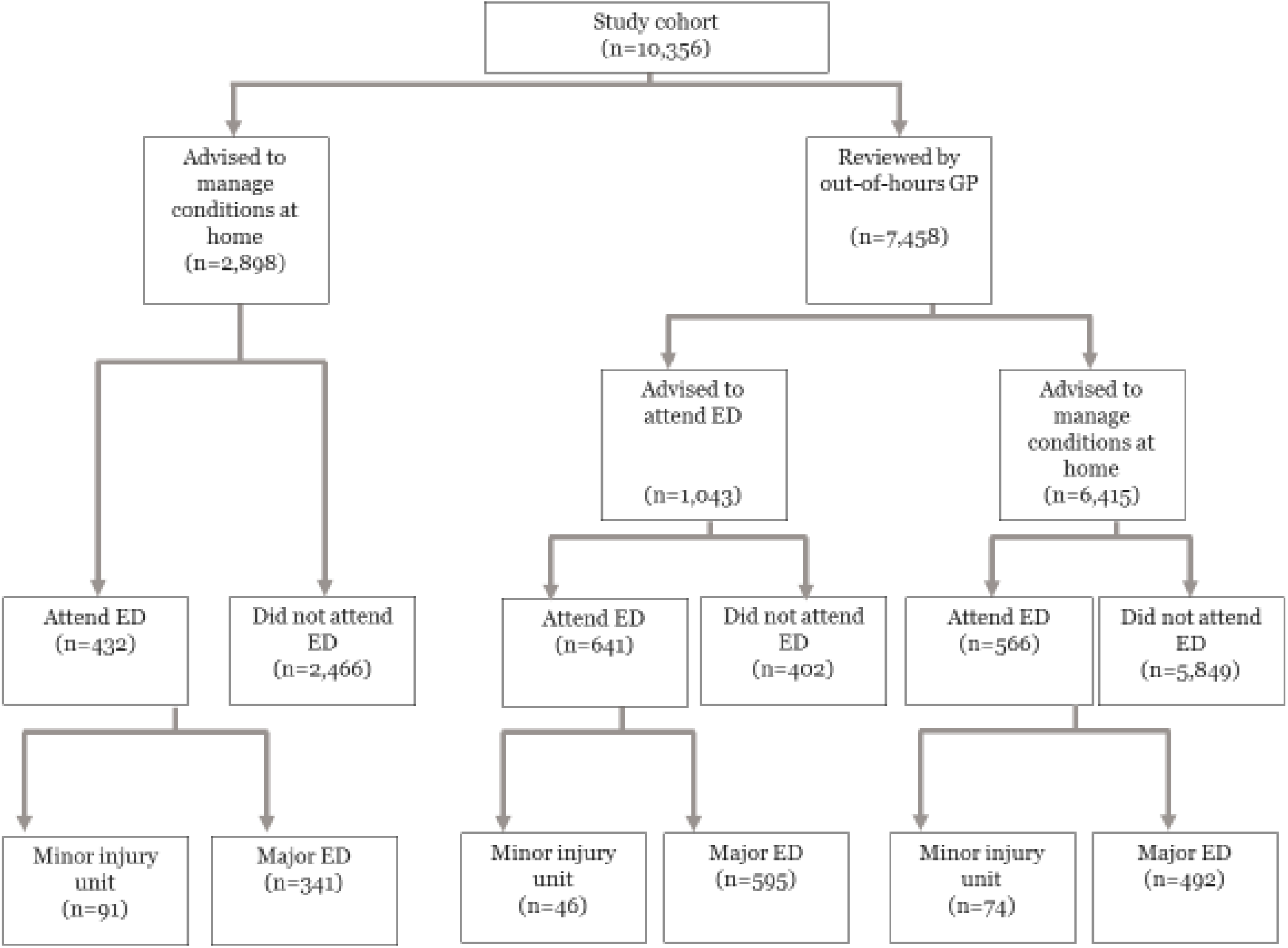
Flow of patient cohort through NHS 111. Shows emergency department utilisation (ED) for the caller cohort. Cohort selection criteria shown in Figure A1. in supplementary material.

**Table 1:** Characteristics of callers (Data show percentage of callers unless stated otherwise)

|  | All calls<br>(n=10,356) | Routed to GP<br>out-of-hours<br>(n=7,458) | NHS 111 call<br>handler advised<br>care at home,<br>without GP<br>review<br>(n=2,898) |
| --- | --- | --- | --- |
| Age of patient in years (mean, SD) | 3.2 (3.7) | 3.1 (3.7) | 3.4 (3.9) |
| Socioeconomic deprivation (1) |  |  |  |
| Quintile 1 (most deprived) | 31.2 | 31.8 | 29.9 |
| Quintile 2 | 33.3 | 33.2 | 33.5 |
| Quintile 3 | 20.3 | 20.0 | 21.0 |
| Quintile 4 | 14.9 | 14.8 | 15.4 |
| Quintile 5 (least deprived) | 0.3 | 0.3 | 0.2 |
| Ethnicity of caller |  |  |  |
| White | 45.9 | 44.8 | 48.8 |
| Mixed/Multiple ethnic group | 10.2 | 10.3 | 9.8 |
| Asian/Asian British | 9.4 | 9.7 | 8.8 |
| Black/African/Caribbean/Black British | 6.4 | 6.6 | 5.8 |
| Other | 9.0 | 8.9 | 9.0 |
| Not stated | 12.4 | 11.9 | 13.5 |
| Missing ethnic group | 6.8 | 7.8 | 4.1 |
| Distance to nearest ED in kilometres (mean, SD) (2) | 1.1 (3.1) | 1.0 (0.5) | 1.3 (5.8) |
| Caller spoke fluent English (3) | 78.4 | 77.7 | 80.3 |
| Patients who were called back by NHS 111 | 15.6 | 11.8 | 25.2 |
| Length of NHS 111 episode in minutes (mean, SD) (4) | 11.8 (14.2) | 10.1 (10.5) | 16.3 (20.2) |
| Caller was 'warm transferred' (5) | 8.9 | 6.0 | 16.4 |
| Call was made during wintertime (November to February) | 43.7 | 44.4 | 42.1 |
| Day of week |  |  |  |
| Weekday (Monday to Friday) | 48.8 | 44.3 | 60.6 |
| Weekend (Saturday or Sunday) | 51.2 | 55.9 | 39.5 |

**Notes**
1 Based on national quintiles of the Index of Multiple Deprivation. Missing data for fewer than 10 calls.
2. Missing data for 693 calls.
3. Missing data for 603 calls.
4. Missing data for 12 calls. 5. Patient was transferred to a clinician whilst on the line. - SD = Standard Deviation.

**Table 2:** Symptoms of callers (Data show percentage of callers)

|  | **All calls**<br>**(n=10,356)** | **Routed to GP**<br>**out-of-hours**<br>**(n=7,458)** | **NHS 111 call**<br>**handler**<br>**advised care at**<br>**home, without**<br>**GP review**<br>**(n=2,898)** |
| --- | --- | --- | --- |
| Breathing difficulty/cough/cold/flu | 11.4 | 13.1 | 7.1 |
| Febrile illness | 5.9 | 7.0 | 3.1 |
| Diarrhoea or vomiting | 10.2 | 11.6 | 6.5 |
| Abdominal pain, constipation or rectal pain | 3.7 | 4.2 | 2.4 |
| Rash or skin problem | 6.5 | 7.0 | 5.2 |
| Injury, limb problem or burn | 3.1 | 2.9 | 3.6 |
| Allergy | 0.7 | 0.8 | 0.4 |
| Ear, nose or throat problem | 5.6 | 6.1 | 4.4 |
| Unwell infant | 15.0 | 17.5 | 8.8 |
| Blisters | 2.0 | 2.3 | 1.0 |
| Chest pain | 0.6 | 0.6 | 0.4 |
| Eye problem | 1.8 | 1.6 | 2.2 |
| Pre-determined plan or specific pathway | 12.2 | 15.1 | 4.8 |
| Requests for health and social care information | 1.3 | 0.9 | 2.1 |
| Dental problems | 8.0 | 8.1 | 7.8 |
| Other | 12.0 | 1.1 | 40.3 |

### Emergency department attendances

Of the 2,898 callers who advised by NHS 111 to manage their health needs at home, and thus were not directed to an out-of-hours GP, 432 callers (14.9%) subsequently attended an emergency department, with 91 callers (3.1%) attending a minor injury unit and 341 callers (11.8%) attending a major emergency department (see Figure 1). In comparison, of the 7,458 callers referred to GP out-of-hours, 1,207 callers (16.2%) subsequently attended an emergency department, with 120 callers (1.6%) attending a minor injury unit and 1,087 callers (14.6%) attending a major department. These crude rates of emergency department utilisation were not statistically different between the group advised to manage their health needs at home at those referred to an out-of-hours GP. For example, Table 3 shows that the unadjusted odds ratio for attendance at any emergency department was 1.10, which was not statistically significant (95% CI, 0.98 to 1.24). Yet, as noted above, the group of patients referred to an out-of-hours GP had more health conditions and were younger than those advised to manage their needs at home. These characteristics that would be expected to lead to higher need for health care.

**Table 3:** Unadjusted and adjusted odds ratios from logistic regression (Selected variables only, n=10,340)

|  | Any emergency department attendance |  | Major emergency department attendance |  | Minor Injury Unit Attendance |  | Emergency admission |  |
| --- | --- | --- | --- | --- | --- | --- | --- | --- |
|  | Unadjusted odds ratio (95% confidence interval) | Adjusted odds ratio<sup>1</sup> (95% confidence interval) | Unadjusted odds ratio (95% confidence interval) | Adjusted odds ratio<sup>1</sup> (95% confidence interval) | Unadjusted odds ratio (95% confidence interval) | Adjusted odds ratio<sup>1</sup> (95% confidence interval) | Unadjusted odds ratio (95% confidence interval) | Adjusted odds ratio<sup>1</sup> (95% confidence interval) |
| Patients who had an out-of-hours GP appointment vs. those who did not | 1.10<br>(0.98-1.24) | 0.86<br>(0.75-0.99) | 1.28<br>(1.12-1.15) | 1.06<br>(0.90-1.24) | 0.47<br>(0.36-0.62) | 0.32<br>(0.23-0.44) | 1.67<br>(1.15-2.42) | 1.29<br>(0.82-2.02) |
| Caller spoke with clinical supervisor ('warm transfer') vs. those who did not | 0.78<br>(0.64-0.95) | 0.77<br>(0.62-0.97) | 0.74<br>(0.56-0.92) | 0.72<br>(0.57-0.92) | 1.13<br>(0.71-1.80) | 1.31<br>(0.75-2.28) | 0.59<br>(0.31-1.11) | 0.76<br>(0.35-1.65) |

Notes
<sup>1</sup> This table shows selected results from the logistic regression model which adjusts for a full range of call and patient characteristics. We report the results for the full set of covariates in the supplementary material.

When controlling for patient characteristics including age and health conditions, callers who were referred to an out-of-hours GP were less likely to attend emergency departments than callers who without an appointment (adjusted odds ratio 0.86, 95% CI, 0.75 – 0.99). When each subset of emergency department type was considered separately, those reviewed by an out-of-hours GP remained less likely to attend minor injury units (adjusted odds ratio 0.32, 95% CI, 0.23 - 0.44), but there was no impact on major emergency department attendance (adjusted odds ratio 1.06, 95% CI, 0.90 – 1.24) – see Table 3.

Callers who spoke with clinical supervisor (‘warm transfer’) while on the line with NHS 111 were less likely to attend emergency departments than other callers (adjusted odds ratio 0.77, 95% CI, 0.62-0.97). When each subset of emergency department type was considered separately, callers who spoke with clinical supervisor remained less likely to attend major emergency department (adjusted odds ratio 0.72, 95% CI, 0.57-0.92), but there was no impact on minor injury unit attendance (adjusted odds ratio 1.31, 95% CI, 0.75-2.28) – see Table 3. Full results reporting the adjusted model results including all covariates are available in Tables A2-A4 in the supplementary material.

### Emergency admissions

Being reviewed by an out-of-hours GP was not associated with lower rates of emergency hospital admission (adjusted odds ratio 1.29, 95 CI%, 0.82 to 2.06) and neither was speaking to a clinical supervisor within the NHS 111 call centre (adjusted odds ratio 0.76, 95 CI%, 0.35 to 1.65) – see Table 3 for selected covariates and Table A5 in for the supplementary material for full model results.

## DISCUSSION

NHS 111 has a significant role in triaging requests for out-of-hours, receiving almost 15 million calls annually. The advice given within NHS 111 telephone calls might therefore have an impact on demand for acute care, and one important factor might be the degree to which callers to NHS 111 are reviewed by clinically-trained staff. We examined a large and representative sample of 10,356 calls to NHS 111 for children and young people in North West London, and found that similar percentages of patients attended emergency departments, irrespective of whether they were advised by NHS 111 to manage their health needs at home or were additionally reviewed by an out-of-hours GP (14.9% vs. 16.2%). Yet, the patients who were reviewed by a GP had more health conditions and were younger on average. After adjusting for these and other patient characteristics, review by an out-of-hours GP was associated with fewer visits to emergency departments (adjusted odds ratio 0.86, 95% CI, 0.75 to 1, p=0.05). In subset analysis by emergency department type we found that those referred to an out-of-hours GP were less likely to attend a minor injury unit, but there was no evidence of an impact on major emergency department attendance.

Our findings suggest that increased clinical input by GPs reduces utilisation of emergency departments following calls to NHS 111 for children and young people. The mechanism behind this finding requires further research. One possibility is that GPs offered treatment or advice that reduced the need for immediate emergency care; another is that callers were more reassured following a consultation with a GP than a non-clinically-trained call handler. In support of the second theory, we note findings from another study that around one-quarter of parents calling the number did not fully have confidence and trust in the first call handler they spoke to at NHS 111.[27] We also note that our finding that reductions in emergency department utilisation were more marked amongst minor injury unit than major departments. It is unknown but plausible that visits to minor injury units are more likely to be prevented through reassurance by a GP than visits to major emergency departments for children and young people Interestingly, we found that although review by a clinical supervisor was associated with similar reductions in emergency department attendances to review by an out-of-hours GP (adjusted odds ratio 0.77, 95% CI, 0.62-0.97), the effect from clinical supervisor review was more pronounced at the major departments. Thus, it is possible that the mechanism of effect was different for clinical supervisor review than GP review.

Importantly, we found that the proportion of patients admitted through the emergency department was similar regardless of whether a patient spoke to a GP or clinical supervisor within the out-of-hours care setting. Therefore, if some emergency department attendances were indeed avoided through clinical input, these were for patients later judged as not requiring hospital admission. Indicating that the complaint did not require acute care, and could be adequately treated within primary care. The findings suggest that there is potential to reduce the use of emergency care, by increasing the degree of clinical input at the beginning of the urgent care pathway.

### Strengths and limitations

Although the major threat to validity in any observational study is confounding, the study was designed to limit its susceptibility to bias. We excluded callers whom NHS 111 directed to the emergency services or advised to attend the emergency department, meaning that we compared two groups of patients: those who were reviewed by a GP and those who were advised to manage their health needs at home. The group of people who were reviewed by a GP were more likely to have a range of health conditions, including breathing difficulties, diarrhoea or vomiting, constipation, and fever. We would expect this group to have higher utilisation of emergency department care. However, despite their higher levels of health need, the group of people who were reviewed by a GP appeared to have similar levels of emergency department attendance even before risk adjustment.

The study benefited from access to the data that was recorded by the NHS 111 health advisors and used when deciding whether or not to refer individual patients to the out-of-hours GP service. Guidance regarding observational studies stresses the need to understand the treatment assignment mechanism and to control for the variables that form part of that decision making process.[28] Thus, from a statistical perspective, this study benefited from the protocolised and computerised approach to decision making, and access to the operational data sets. However, it must be acknowledged that confounding remains a limitation. Ethnographic studies of the use of computerised decision support systems in urgent and emergency care have identified a number of ways in which the human and technological dimensions interact to produce a sociotechnical system, with various forms of discretion existing regarding the ultimate dispositions.[29] For example, patients might be referred to a GP when they seem in obvious distress, regardless of the answers to the questions posed by the NHS Pathways algorithm. Therefore, notwithstanding the use of operational data, it is possible that the two groups of patients differed in ways that we could not observe, and this would mean that our adjusted odds ratios are biased. The most likely direction of the bias is towards the null – in other words, we have probably underestimated, rather than overestimated, the effect of GP review on reducing emergency department attendances. Thus, we cannot entirely discount the possibility that GP review reduced attendances at the major departments as well as at the minor injury units, but remain confident about our conclusion that the overall level of emergency department utilisation across both minor and major departments was lower following GP review than would be the case if the same patients had been advised by NHS 111 call handlers to manage their health conditions at home. The availability of GPs to review patients following calls to NHS 111 therefore seems to aid efforts to reduce emergency department use.

Following this study, it is natural to ask whether levels of GP (and indeed, clinical supervisor) input should be expanded, either directly within NHS 111 or by establishing models that allow NHS 111 calls to be referred to GPs working in other services. In considering this possibility, we urge some caution. Our results do indeed suggest that GP input may reduce demand for emergency care, but one limitation of our study is that it related to three areas of North West London, which is more socioeconomically deprived than England as a whole (Table 1) but might have a higher degree of integration between NHS 111 and out-of-hours GP services than other areas, since in North West London a GP-led not-for-profit organisation (the London Central & West Unscheduled Care Collaborative) is responsible for both services. These features may explain some of our findings, such as the high referral rate observed from NHS 111 to GP out-of-hours services. It is likely that the impact of GP review is different in other settings. Furthermore, the impact of GP review may differ for other patient groups than those who were referred to GPs in this particular study (for example, adult patients). A consideration of the impact of GP review should also bear in mind outcomes other than attendance at emergency departments, such as patient experience and safety.[30] These cannot be assumed to correlate with service utilisation. Finally, there may be unintended impacts of increasing GP input in NHS 111 calls, given that the number of GPs is limited.

### Conclusions

There remains a need to manage the demand for urgent and emergency care. In England, the number of attendances at emergency departments increased from 14.0 to 22.4 million between 2003 and 2014.[31] Around 20% of attendances were for children aged under 15,[32] and a notable minority of paediatric attendances are for conditions or symptoms that could have been safely managed outside of secondary care.[33,34]

In our study children and young people who received an appointment with GP out-of-hours care following a call to NHS 111, or had input during the call from a clinical supervisor were less likely to attend emergency departments within 10 hours of the call. There was no impact demonstrated on subsequent emergency hospital admissions. Therefore, clinical input during the NHS 111 call, and integration with a GP out-of-hours service may reduce inappropriate emergency department visits by children and young people.

## Acknowledgements

We would like to thank Andreas Magusin at North and East London Commissioning Support Unit for extracting the data required for this study.

## Contributorship

AS, IM and DH conceived the study. AS and AR came up with the statistical analysis plan. AR, RP, CR carried out the analysis. SD, AR and AS interpreted the analysis, drafted and finalised the paper. All authors approved the final draft of the paper.

## Transparency declaration

The senior author* affirms that this manuscript is an honest, accurate, and transparent account of the study being reported; that no important aspects of the study have been omitted; and that any discrepancies from the study as planned (and, if relevant, registered) have been explained.

*The manuscript’s guarantor.

